# Mannoprotein Cig1 Contributes to the Immunogenicity of a Heat-Killed F-box Protein Fbp1 *Cryptococcus neoformans* Vaccine Model

**DOI:** 10.1101/2025.06.17.660137

**Authors:** Samantha Avina, Siddhi Pawar, Amariliz Rivera, Chaoyang Xue

**Affiliations:** School of Graduate Studies at the Health Science Campus, New Jersey Medical School, Rutgers University Newark, New Jersey, USA; Public Health Research Institute, and Department of Microbiology, Biochemistry, and Molecular Genetics, New Jersey Medical School, Rutgers University, Newark, New Jersey, USA; Center for Immunity and Inflammation, and Department of Pediatrics New Jersey Medical School, Rutgers University Newark, New Jersey, USA

**Keywords:** HK-*fbp1*, Fungal vaccine, Mannoprotein, Cig1, murine model, immunogenicity

## Abstract

Currently, no fungal vaccine exists for clinical use while fungal infections are responsible for over 1.5 million deaths every year. Our previous studies identified a *Cryptococcus neoformans* mutant strain *fbp1*Δ as a potential vaccine candidate. This mutant strain contains a deletion of the F-box protein Fbp1, a key subunit of the SCF E3 ligase complex necessary for ubiquitin-mediated proteolysis. Vaccination with heat-killed *fbp1*Δ (HK-*fbp1)* can elicit protection against *C. neoformans* parental strain and its sibling species *C. gattii* in an interferon gamma (IFN-γ) dependent Type 1 immune response. However, we have yet to decipher the immunogenic factor(s) expressed by the *fbp1*Δ mutant that are responsible for the induction of the protective immune response. In this study, we have identified that capsule plays an important role in HK-*fbp1* vaccine mediated protection, as acapsular HK-*fbp1* cells showed diminished protection against wild type challenge. Additionally, our studies have shown that Cytokine Inducing Glycoprotein 1 (Cig1), a GPI anchored mannoprotein, is regulated by Fbp1 and contributes to the immunogenicity of HK-*fbp1*. Deletion of Cig1 in the *fbp1*Δ background resulted in decreased recruitment of anti-fungal effector T cells and diminished production of protective inflammatory cytokines by the host. Furthermore, loss of Cig1 in the *fbp1*Δ mutant resulted in reduced protection in vaccination survival studies at lower vaccine inoculum doses compared to HK-*fbp1*. In aggregate, these findings demonstrate Cig1 is an antigen contributing to the immunogenicity of HK-*fbp1* that may be utilized to further optimize the HK-*fbp1* fungal vaccine as a tool in the arsenal against invasive fungal infections.

## Introduction

*Cryptococcus neoformans* is an encapsulated yeast that is the primary causative agent of cryptococcosis, that predominantly affects immunocompromised populations. *C. neoformans* has been identified as the leading fungal pathogen responsible for fungal encephalitis and is responsible for ∼20% of AIDS related deaths worldwide^1^. In 2022, the inaugural World Health Organization fungal priority pathogens list ranked *C. neoformans* in the top research critical priority group of fungal pathogens as cases of invasive fungal disease continue to rise^2^. External factors that contribute to increased susceptibility of *C. neoformans* include increased use of immunosuppressants (i.e., cancer chemotherapies, organ transplant recipients) and potentially climate change as global temperatures rise^3,4^. Invasive fungal pathogens like *C. neoformans* are eukaryotes that share large similarities of cellular machinery with human hosts, making treatment options limited, prolonged, and often unsuccessful. As *C. neoformans* continues to be a rising global health concern, the greatest limit to the anti-fungal arsenal is lack of anti-fungal vaccination strategies. Currently, no fungal vaccines are available for clinical use.

Protection against *C. neoformans* is primarily driven in a CD4^+^ T cell dependent manner. HIV/AIDS immunocompromised populations deficient in CD4^+^ T are highly susceptible to cryptococcosis disease and thus emphasize the importance of T cell mediated immunity in the context of host response against cryptococcal infection^5^. Humoral immunity is also understood to contribute to host immunity as previous studies have shown antibodies generated against capsule components GXM and Gal-XM ameliorate fungal burden when administered to murine models but not sufficient in conferring protection against *C. neoformans* challenge in murine models^6–8^. More importantly, a balance of inducing a T_H_1 protective inflammatory response while not generating a detrimental over inflammatory response is key in protection of different vaccine strain candidates. In multiple *C. neoformans* based vaccine candidates, the ability to elicit production of IFN-γ by the host CD4^+^ T or CD8^+^ T cells in the absence of CD4^+^ T cells upon vaccination and subsequent challenge has been critical^9–16^.

In our previous studies we have identified a *C. neoformans* mutant strain lacking F box protein 1 (*fbp1*Δ) as a potential vaccine candidate. The F-box protein is a part of the E3 ubiquitin ligase complex responsible for ubiquitin-mediated proteolysis of target proteins by the 26S proteosome in eukaryotic systems that are required for proper cellular functions. It has been established that while Fbp1 is not required for development of classical virulence factors (melanin, chitin/chitosan, capsule size, etc.)^17^, it does regulate cell size, sexual reproduction, and is essential for fungal virulence^18–20^. Interestingly, the *fbp1Δ* mutant infection triggered strong protective Th1 immunity, and this protective immunity remains even in animals immunized with the heat-killed fbp1 cells (HK-*fbp1*). Therefore HK-*fbp1* has been developed as an excellent vaccine candidate. However, the immunogens in HK-*fbp1* vaccine remain uncharacterized.

Cryptococcal mannoproteins are heavily glycosylated and reside in the capsule closer to cell wall^21,22^ that are highly immunogenic antigens and can stimulate T_H_1 response of CD4^+^ T cells to produce cytokines including IL-6, IL-10, IFN-γ, and TNF-α^23–25^. Mannosylation of cryptococcal mannoproteins also enhances their immunogenicity for mannose receptors for uptake and presentation to T cells ^26–28^. Mannoproteins MP-98 and MP-88 have been described to reside close to the cell wall rather than as integral parts of the capsule and largely contribute as immunogens that can be secreted from the cell to trigger immunogenic responses^21,24,29,30^. Mannoprotein deficient strains including the chitin deacetylase triple mutant (*cda1*Δ *cda2*Δ *cda3*Δ) has been shown to be critical to enhancing its immunogenicity and utilized as a promising *C. neoformans* vaccine candidate^14,31–34^. Yet, studies have also found that cryptococcal mannoproteins, like Krp1, contribute to capsule structuring mainly by subtle altering distribution of glycans in the cell wall^35^. Whether mannoproteins play a role in the immunogenicity of all current cryptococcal based fungal vaccine candidates has yet to be thoroughly explored.

In this study we set out to investigate the role of mannoproteins and capsule in mediating the immunogenicity of the heat killed *fbp1*Δ (HK-f*bp1)* vaccine protection. Here we report that capsule plays an important role in HK-*fbp1* vaccine mediated protection, as acapsular HK-*fbp1* cells showed diminished protection against wild type challenge. Additionally, serum from HK-*fbp1* vaccinated mice showed to bind to distinct regions of the capsule in *fbp1Δ* strains versus parental H99 strain indicating an importance of immunogenic antigens in the capsule specific to *fbp1*Δ. We found that Cytokine Inducing Glycoprotein 1 (Cig1, CNAG_01653), a GPI anchored mannoprotein with a PEST domain sequence, is upregulated in *fbp1*Δ and elicits antibody production in HK-*fbp1* vaccination sera. Furthermore, we found that loss of Cig1 in the *fbp1*Δ background diminishes the protective host response upon vaccination and induction of T_H_1 in live strain infection. While differences in protection were only observed at lower vaccine inoculum doses, these studies suggest Cig1 is an immunogenic antigen upregulated in *fbp1*Δ that contributes to the enhanced immunogenicity of the HK-*fbp1* vaccine and may be a substrate regulated by *fbp1*Δ.

## Results

### 1.1 Capsule production contributes to HK-fbp1 vaccine protection against H99 Cryptococcus challenge

To determine what antigens contribute to the immunogenicity of *fbp1*Δ, we first examined the role of capsule as it is the first cellular surface that encounters the host and masks cell wall antigens. In previous studies, we broadly screened for differences in classical virulence factors of *fbp1*Δ and saw no differences in capsule size or secreted GXM^17^. In TEM imaging comparing H99 and *fbp1*Δ, we also saw no major architectural changes between the strains (**Figure 1C**). To ascertain whether there are specific regions of the capsule that are immunogenic, we utilized HK-*fbp1* vaccinated mouse serum in an immunofluorescence assay to visualize and compare localization of HK-*fbp1* vaccine antibody binding. Interestingly, there was a difference in HK-*fbp1* serum binding to the capsule of *fbp1*Δ compared to H99 (**Figure 1A**). Specifically, serum from HK-*fbp1* vaccinated mice bound to the outer rim of capsule induced *fbp1*Δ compared to H99 and a higher percentage of *fbp1*Δ cells showing serum binding localized to the outer region of the capsule (**Figure 1B**). While there was an observed difference in localization, antibodies still bound robustly to the acapsular cap59 strain controls (**Figure 1A**). Due to previous exposure to HK-*fbp1* Cryptococcal antigens, we anticipated serum from HK-*fbp1* vaccinated mice would produce an array of antibodies to many regions of the Cryptococcal cell. As antibodies from HK-*fbp1* serum bound distinctively to acapsular strains we desired to confirm whether capsule is required for HK-*fbp1* protection.

**Figure 1.**
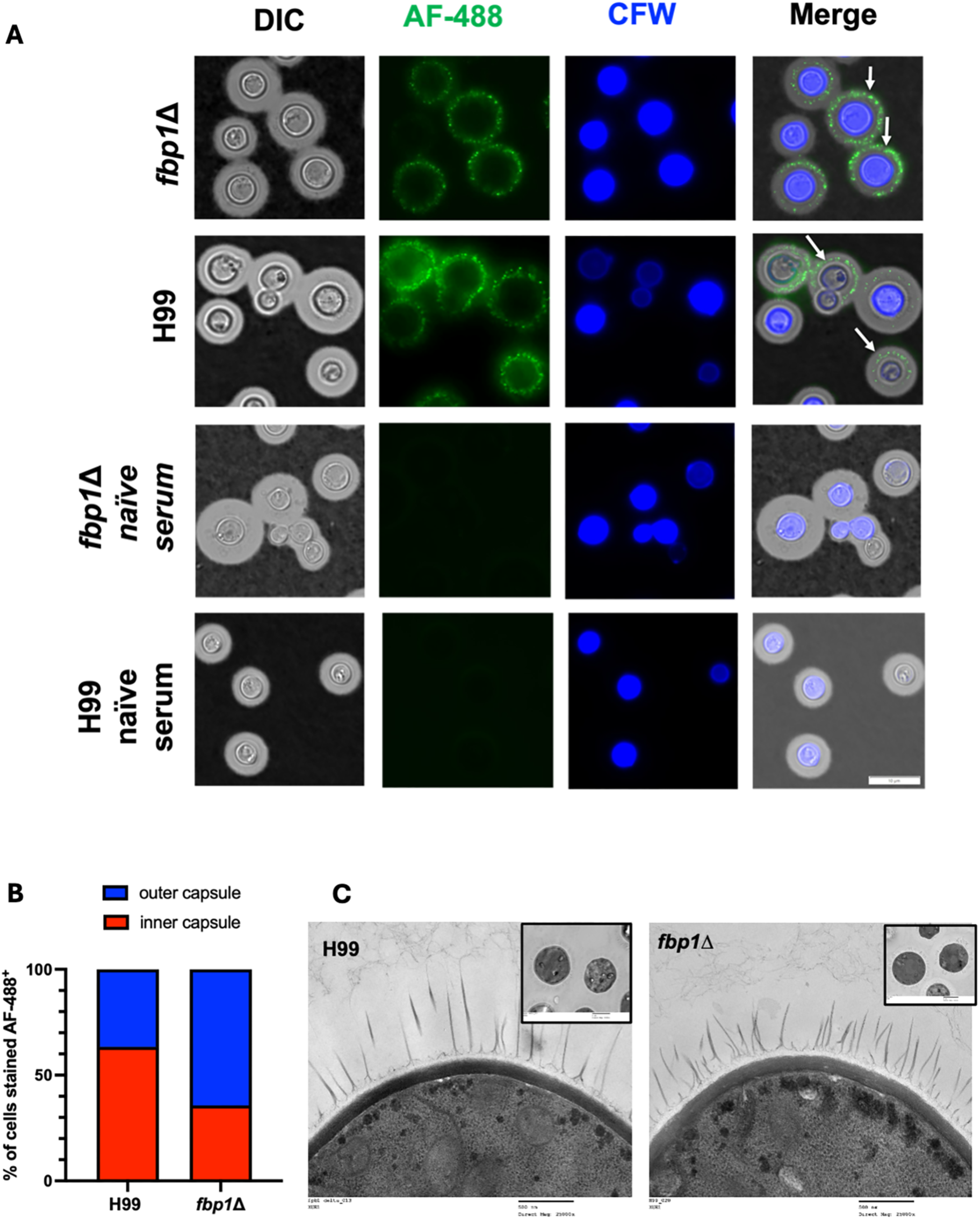
HK-fbp1 vaccinated mouse antibodies bind to different regions of *fbp1*Δ *Cn* capsule. **A.** Wildtype H99 and *fbp1*Δ grown in minimal media capsule inducing media for 3 days and incubated with HK-*fbp1* vaccinated mouse serum conjugated to AF-488. Cells were imaged in India ink to show capsule and stained with calcofluor white to contrast fungal cell wall at 100x magnification. Cells were also incubated with serum from naïve mice as control. Images taken in DIC GFP and DAPI channels were overlayed and merged to show distinction of binding by antibodies in vaccinated serum. **B.** Quantitative ratio of binding on the inner capsule versus outer capsule in percentage of total cells counted as AF-488+. **C.** TEM imaging of H99 and *fbp1*Δ comparing cell wall and capsule structures. Bar=500nm.

We next used an acapsular *fbp1*Δ strain (*cap59*Δ *fbp1*Δ) to investigate the adaptive immune response in the context of vaccination compared to *fbp1*Δ. In this vaccination experiment utilizing the previously established intranasal vaccination method (**Figure 2A**) to compare heat-killed acapsular strains HK-*cap59 fbp1* and HK*-cap59*, we observed that mice vaccinated with either heat-killed acapsular strain showed similar CD4^+^ and CD8^+^ T cell population percentages in the bronchiolar lavage fluid (BALF) (**Figure 2B**). However, heat-killed acapsular strain vaccinated cohorts showed diminished recruitment of CD4^+^ T cells in the lung compared to capsular HK-*fbp1* (**Figure 2C**). Furthermore, the functionality of CD4^+^ cells from the BALF to produce key protective cytokine IFNγ was also diminished as observed via flow cytometry (**Figure 2C and Figure S1**). Survival studies showed that heat-killed acapsular strain vaccinated cohorts showed significant decrease in survival compared to the capsular HK-*fbp1* and succumbed to lethal cryptococcal meningitis (**Figure 2E-2F**). These findings supported that while capsule is required for full HK-*fbp1* mediated protection, it is not sufficient as 40% of the acapsular HK-*fbp1* strain vaccinated cohorts still survived until the termination of the survival (endpoint) at day 70 post-challenge. In the remaining survivors, fungal dissemination in the lung and brain was present, indicating these cohorts may have eventually succumbed to cryptococcal infection. These results suggest that capsule is required but not sufficient in inducing HK-*fbp1* mediated immune protection. Thus, we proceeded to assessing the role of non GXM or GalXM based antigens.

**Figure 2.**
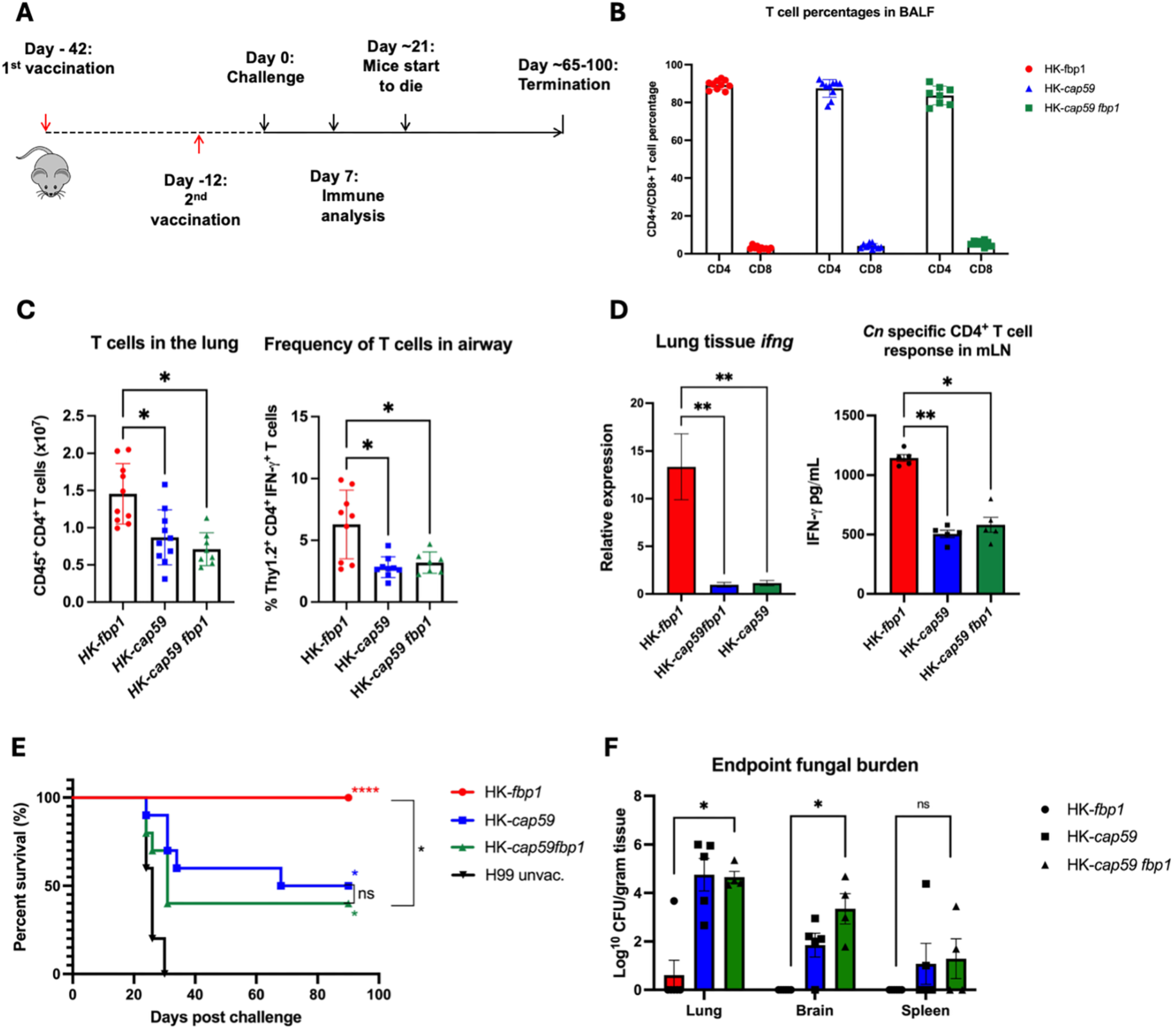
Capsule is required for full HK-*fbp1* vaccine mediated protection. **A.** Schematic of vaccination strategy. Mice are vaccinated with HK-*fbp1* or acapsular strains (HK-*cap59* or HK-*cap59 fbp1*) 42 days and 12 days before challenge with 1×10^4^ H99 wildtype strain. **(B-D)** Day 7 post H99 challenge immune response. IFN-γ producing CD4^+^ T cells in BALF (**B**), IFN-γ cytokine readout from isolated T-cells of vaccinated mice presented H99 antigen (**C**), qPCR from total lung tissue, and ratio of CD4^+^ to CD8^+^ T cells in the vaccinated cohorts (**D**). * p<0.05, **p<0.001, ***p<0.0005, ****p<0.0001, Kruskal-Wallis test with Dunn’s multiple comparison test. **(E-F)** Survival curve of mice vaccinated with HK-*fbp1* or acapsular strains (**E**) and fungal burden (**F**) of remaining survivors. ***p<0.0005, ****p<0.0001 (log rank [Mantel-Cox] test). Data is representative of two independent experiments.

### 1.2 Cytokine Inducing Glycoprotein 1 (CIG1) is overexpressed in fbp1Δ and a substrate of Fbp1 localized to the cell wall

Because mannoproteins are known cell surface immunogens important for host immune recognition^21,23,36^, and we did not detect any difference in chitin and chitosan levels in the *fbp1*Δ cells, we decided to investigate the production of mannoproteins in H99 and its *fbp1*Δ cells by Concanvalin A (ConA) staining. We found that *fbp1*Δ cells showed higher FITC-ConA signal intensity in comparison to H99 (**Figure 3A),** suggesting an overproduction of mannoproteins in the mutant. To identify the genes responsible for the mannoprotein difference, we screened eight reported mannoprotein encoding genes using qRT-PCR under YPD growth condition. Our data demonstrated that one of them, Cytokine Inducing Glycoprotein 1 (*CIG1*, CNAG_01653), was highly upregulated in the *fbp1*Δ background (**Figure 3B**). Interestingly, in our previous Fbp1 pull down studies using a ligase-trapping system, we also identified Cig1 as one of Fbp1 interacting proteins as potential SCF(Fbp1) substrates (unpublished). We posit that Cig1 may be a substrate of Fbp1 that can undergo post-translational modifications, at the same time it is also transcriptionally regulated by Fbp1, either directly or by modulation of other downstream substrates. These studies suggest that Cig1 may be an immunogen overproduced in *fbp1*Δ cells. Therefore, we decided to further analyze the potential role of Cig1 in *fbp1*Δ mediated immune activation. Cig1 is a heavily glycosylated GPI anchored mannoprotein (**Figure 3C**). It has been identified as a hemophore required for iron nutrient sensing and uptake for *C. neoformans* in host conditions ^37^. Excitingly, while we were analyzing the Cig1 regulation by Fbp1 E3 ligase, it was recently independently reported that Cig1 is an antigen in *C. neoformans* Znf2^OE^ vaccine that can be utilized for mRNA-based vaccine development ^38^. This independent study further supported our hypothesis that Cig1 may be an important immunogen in the HK-fbp1 vaccine model. To confirm that Cig1 interacts with Fbp1, we generated a C-terminus mCherry tagged *CIG1* construct under the control of the *C. neoformans* actin promotor (*P_ACT1_-CIG1:mCherry*) and transformed it into a *C. neoformans* strain expressing Fbp1:FLAG (**Figure 3C**). The expression of Cig1 was confirmed by mCherry expression. We found that the mCherry signal was primary localized to the plasma membrane, consistent with its function as a GPI-anchored mannoprotein (**Figure 3C**). We conducted a Co-immunoprecipitation (Co-IP) experiment using this *C. neoformans* strain expressing both Fbp1:FLAG and Cig1:mCherry proteins and confirmed the interaction between Fbp1 and Cig1 (**Figure 3D**). Cig1 has a PEST domain, a putative signature of short half-life proteins often subjected to E3 ligase-mediated ubiquitination, further indicating that Cig1 is likely a substrate of the Fbp1 and subjected for Fbp1-mediated ubiquitination and degradation. Indeed, our in vivo ubiquitination assay revealed that Cig1 is extensively ubiquitinated (**Figure 3E**).

**Figure 3:**
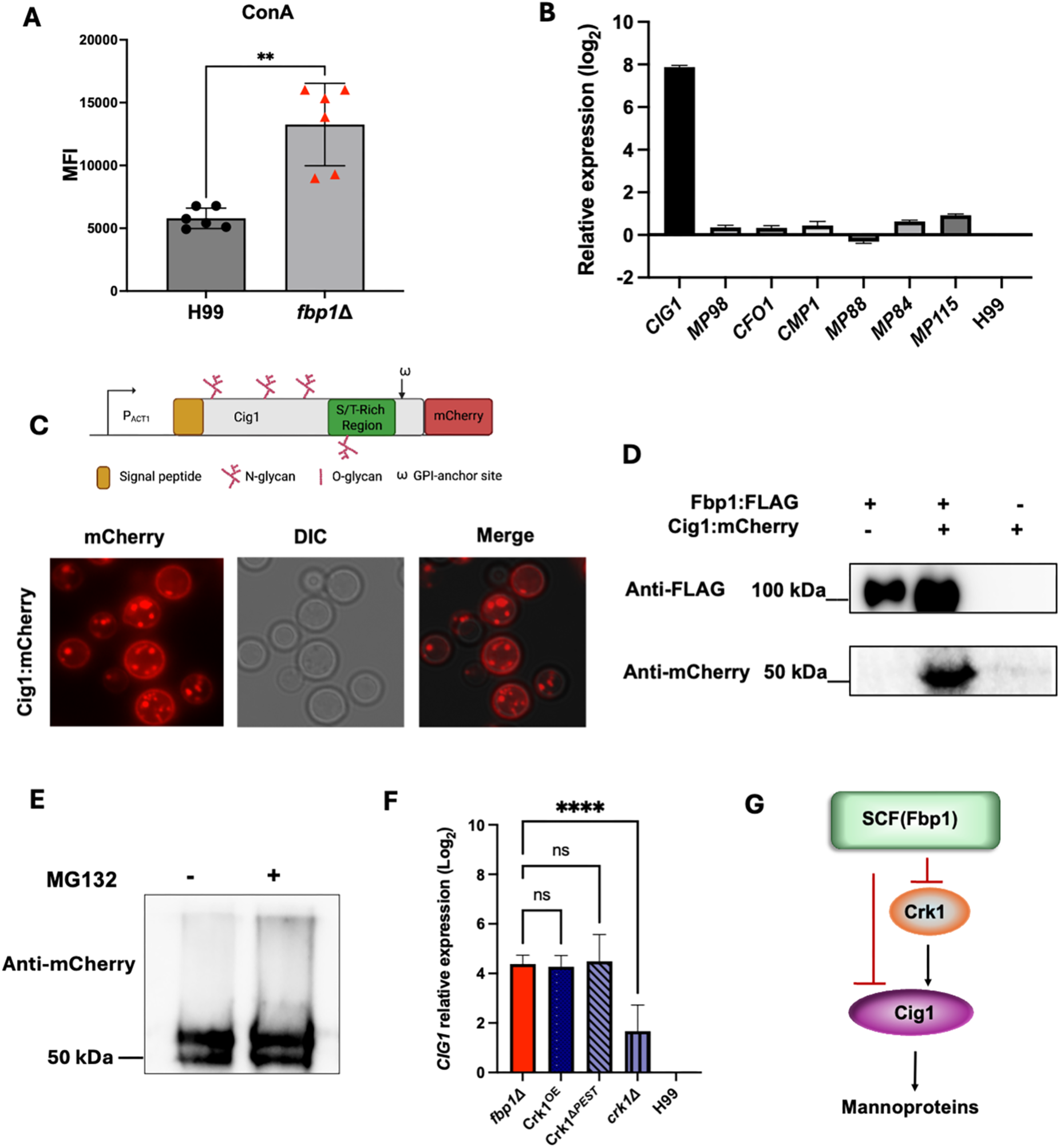
Cig1 interacts with Fbp1 and is overexpressed at the transcriptional level in *fbp1Δ.* **A.** Comparison of MFI ConA-FITC positive stained cells amongst mutant strains. **p<0.0005, non-parametric Mann Whitney test. Data is cumulative of two independent experiments. **B.** Expression of *CIG1, CMP1, CFO1, MP88, MP84, MP98, MP115* and *GAPDH* analyzed by qRT-PCR assay. Gene expression levels were normalized using the endogenous control gene *GAPDH* and analyzed using ΔΔCT methods. **C.** Diagram of Cig1:mCherry fusion construct generation and Cig1 protein **glycosylation sites** N-glycosylation sites (red chains), PEST domain predicted region (green box), and a transmembrane domain region (ω). Cig1 is fluorescently tagged with an mCherry protein and localized to the cell membrane. **D.** Co-immunoprecipitation of Fbp1:FLAG with Cig1:mCherry from strain lysates expressing only Fbp1:FLAG, Cig1:mCherry, or both Fbp1:FLAG and Cig1:mCherry. Immunoprecipitated samples were immunoblotted with either anti-FLAG (top) or anti-mCherry (bottom). **E**. In vivo ubiquitination assay to confirm Cig1 is a substrate of Fbp1 E3 ligase. Total proteins from H99 strain with or without proteosome inhibitor MG-132 were used for the assay. **F.** Expression of *CIG1* in multiple listed strain backgrounds analyzed by qRT-PCR assay. Gene expression levels were normalized using the endogenous control gene *GAPDH* and analyzed using ΔΔCT method. *Statistical value. **G.** A proposed model of Cig1 regulation by the SCF(Fbp1) E3 ligase.

These studies confirmed that Cig1 is a substrate of Fbp1 that can undergo post-translational modifications. However, it did not address how Fbp1 also transcriptionally regulates *CIG1* expression. In our previous studies we identified that Crk1, a CDK related kinase 1, is a direct downstream substrate of Fbp1^18^. To test the hypothesis that Fbp1 may regulate *CIG1* expression through its regulation of Crk1 protein, we measured the *CIG1* expression in wild type, *crk1*Δ and CRK1 overexpression (*CRK1^OE^*) allele expressing strains. Our qRT-PCR assay showed that *CIG1* expression is significantly induced in *CRK1* overexpression strains, including *CRK1*^Δ^*^PEST^* and *CRK1^OE^* backgrounds, at a level similar to *fbp1*Δ, and reduced in the *crk1*Δ background, indicating that *CIG1* expression is positively regulated by Crk1 (**Figure 3F**). Hence, our data support the model that Fbp1 regulates Cig1 at two different levels. Namely, Fbp1 suppresses *CIG1* gene expression by degrading its substrate Crk1, then Fbp1-mediated Cig1 ubiquitination also promotes its degradation (**Figure 3G**). Such dual level regulation also suggests a tight functional association between Fbp1 and Cig1 in modulating cellular function, leading to the significant overproduction of Cig1 in the *fbp1*Δ background.

### 1.3 Cig1 is required for fungal cell wall integrity and macrophage interaction

Consistent with previous findings, the *cig1*Δ single mutant did not show any clear phenotypic defect compared to wild type. We then constructed an *fbp1*Δ *cig1*Δ double mutant to characterize the phenotype of *fbp1*Δ in the context of Cig1 deletion. Classical phenotypic characterization of the *fbp1*Δ *cig1*Δ mutant strains revealed only modest phenotypic differences compared to the parental *fbp1*Δ on cell wall and membrane integrity (**Figure S2**). The *fbp1*Δ *cig1*Δ strain showed growth inhibition on Congo Red and L-DOPA agar media when incubated at 37°C. No inhibition was observed when the same plates were incubated at 30°C. In an *in vitro* phagocytosis assay using macrophage-like cell line J774, the *fbp1*Δ *cig1*Δ cells showed a decrease in phagocytosis when compared to the *fbp1*Δ single mutant, to a level similar to H99 (**Figure 4A**). Furthermore, the *cig1*Δ mutant also showed low percentages of phagocytosis resembling wildtype H99 compared to *fbp1*Δ. These *in vitro* assays and phenotypic characterizations show that Cig1 moderately alters *fbp1*Δ growth in different stress conditions (i.e., cell wall integrity and melanin production) and initial interactions with host immune cells.

**Figure 4:**
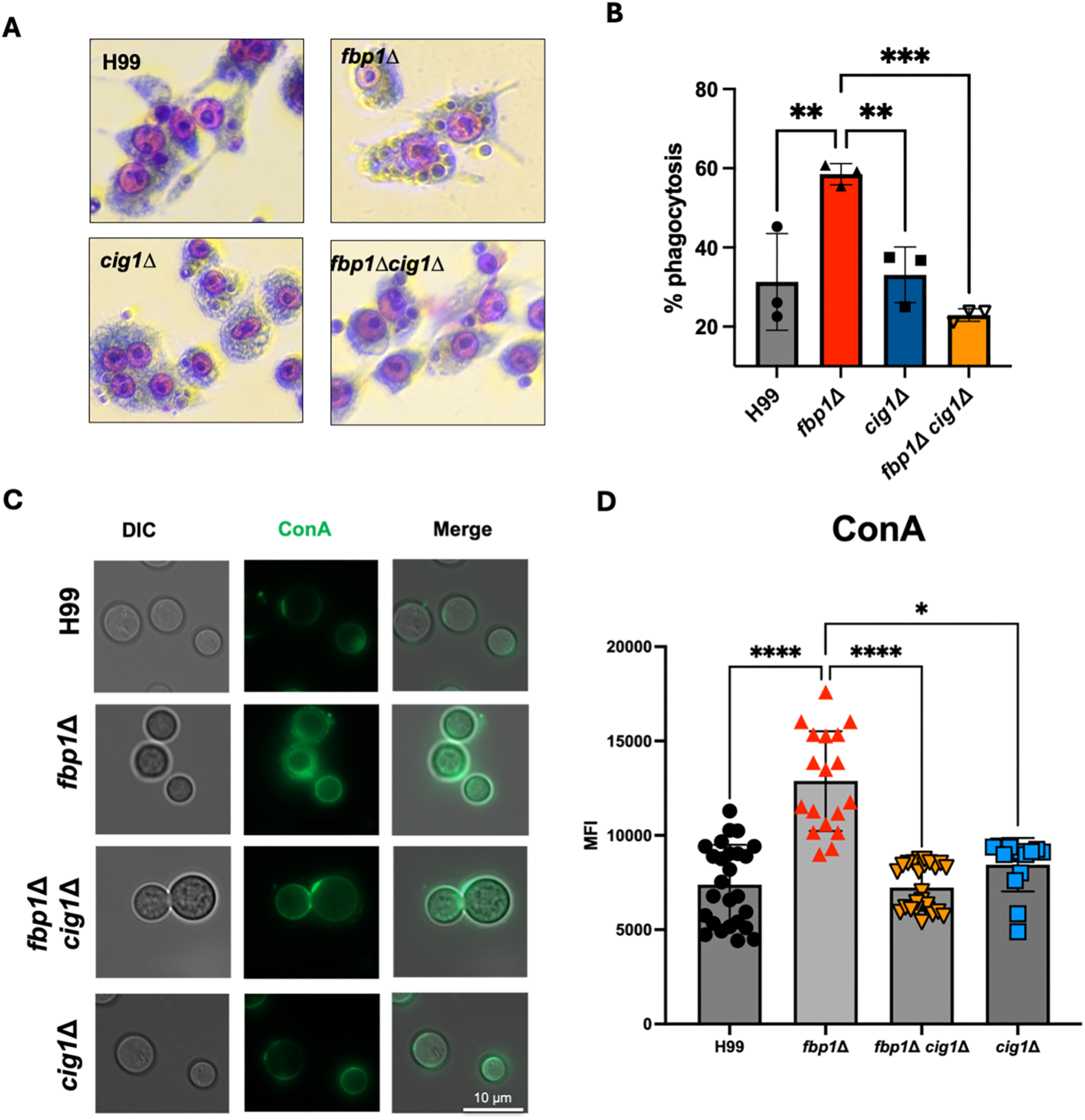
Cig1 mannoprotein alters mannose presence in cell wall and phagocytosis in *fbp1*Δ. **background. A.** 5×10^4^ activated Raw26 MΦ cells incubated with 2×10^5^ opsonized *C.n* for 2 hr. Percentage of phagocytosis of *C.n* with representative images shown at 20x magnification. B. Qualification data of the percentage of yeast cell engulfed by macrophages to determine the phagocytosis efficiency * p<0.05, **p<0.005, ***p<0.0005, One-way ANOVA with Dunnett’s multiple comparisons test. Data is representative of two independent experiments. **C.** Comparison of MFI ConA-FITC positive stained cells amongst mutant strains. **D.** Quantification of ConA staining signal for each listed strain. *p<0.05, ****p<0.0001, Kruskal-Wallis test with Dunn’s multiple comparison test. Data is cumulative of six independent experiments.

To confirm that loss of Cig1 indeed alters *fbp1*Δ mannose levels, we stained all mutants with ConA and compared mutants via flow cytometry analysis of ConA positive stained cells. This experiment showed that *fbp1*Δ did have higher mannose levels compared to H99 and the *cig1*Δ mutant (**Figure 4B**). Furthermore, the loss of high ConA positive signal in the *fbp1*Δ *cig1*Δ double mutant revealed Cig1 is a contributor to high mannose presence in the *fbp1*Δ mutant.

### 1.4 *Loss of Cig1 in fbp1*Δ diminishes induction of protective T_H_1/T_H_17 immune response

To determine the role of Cig1 in *fbp1*Δ-mediated host immune response during live infection, we infected murine cohorts comparing *fbp1*Δ and *fbp1*Δ *cig1*Δ to analyze T cell populations and function (**Figure 5A**). Both mutant strain infected cohorts were administered higher inoculums (10^6^ CFU/mouse) than H99 and *cig1*Δ (10^5^ CFU/mouse) controls to reach comparable fungal burden in the infected lungs for immunological comparison as the *fbp1*Δ parental strain has significantly reduced lung CFU after 3 day post-inoculation as described in our previous publications^17^. Compared to *fbp1*Δ, the mice infected with *fbp1*Δ *cig1*Δ cells had diminished CD4^+^ T cell recruitment in the lung and significantly reduced production of key protective cytokines (i.e., IFNγ, IL-17A, and TNFα) in the airway (**Figure 5B-5C**). To further confirm these functional differences, we performed complementary CD4^+^ T cell recall experiments utilizing primed T cells from the mediastinal lymph nodes (mLN) presented with H99 antigen and measure cytokine production differences. Similar to our flow cytometric data, *fbp1*Δ *cig1*Δ infected animals produced lower key protective cytokines compared to *fbp1*Δ (**Figure 5D**). However, when assessing fungal burden differences 7 days post infection, both *fbp1*Δ and *fbp1*Δ *cig1*Δ mutant strains had low fungal burden compared to H99 and *cig1*Δ (**Figure 5E**). These findings suggest loss of Cig1 altered the host immunological response to *fbp1*Δ infection but does not change the avirulence of *fbp1*Δ that inhibits fungal dissemination.

**Figure 5:**
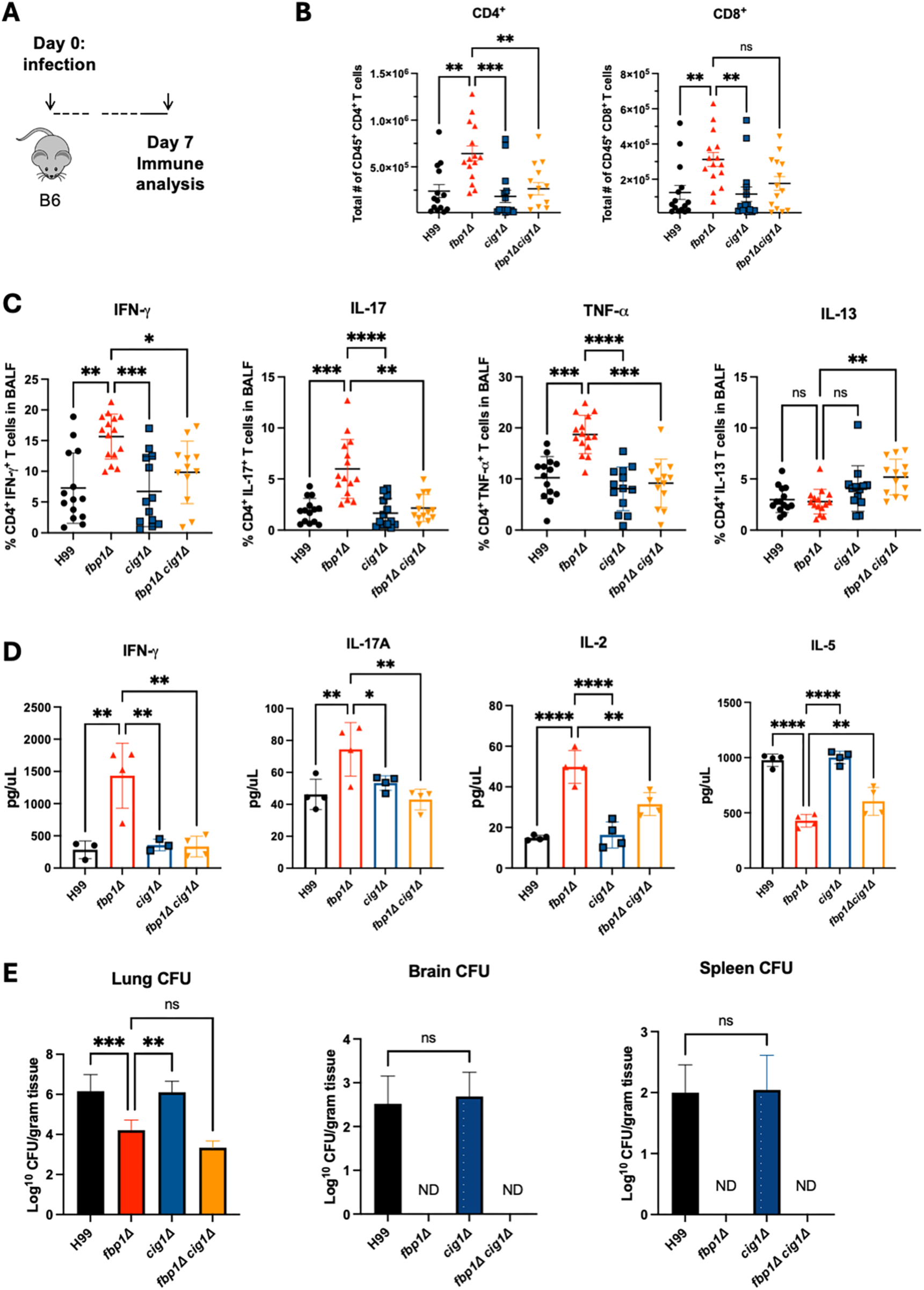
Loss of Cig1 in *fbp1*Δ background diminishes T_H_1 and T_H_17 response at day 7 post infection compared to parental *fbp1*Δ strain. **A.** Schematic of infection with live strains at 10^6^ (*fbp1*Δ, *fbp1Δ cig1*Δ) and 10^5^ (H99, *cig1*Δ) in B6 mice (n=4 or 5 per group). **B.** CD4^+^ and CD8^+^ T cell populations in lung at day 7 post infection. **C.** Cytokine readout from CD4^+^ T cells in BALF 7 days post infection. Kruskal-Wallis test with Dunn’s multiple comparison test was used for both B and C data sets. * p<0.05, **p<0.001, ***p<0.0005, ****p<0.0001. **D.** Cytokine readout from mLN purified CD4^+^ T cells stimulated with H99 antigen for 48-72 hr. **E.** Fungal CFUs for lung, brain, and spleen of mice day 7 post infection. One-way ANOVA with Dunnett’s multiple comparisons test was used for data analysis in **D** and E. * p<0.05, **p<0.005, ***p<0.0005. Data is representative of three independent experiments.

### 1.5 HK-fbp1 primed CD4^+^ T cells elicit protective cytokine responses to Cig1 isolated antigen and contribute to HK-fbp1 vaccination-mediated host protection

To further dissect the importance of Cig1 in *fbp1*Δ immunogenicity, we vaccinated murine cohorts with HK-*fbp1* and assessed whether HK-*fbp1* primed CD4^+^ T cells responded to purified Cig1 antigen (**Figure 6A and SF3**). HK-*fbp1* primed single cell suspension lung tissue samples were restimulated with either whole cell H99 sonicated antigen, purified Cig1 antigen, or no antigen ex vivo and assessed for IFN-γ production by activated CD4^+^ T cells via flow cytometry (**Figure SF4**). Results showed that HK-*fbp1* vaccinated CD4^+^ T cells in the lung environment did produce a protective IFN-γ cytokine response to Cig1 antigen compared to the no antigen controls (**Figure 6B**). Furthermore, sonicated H99 antigen stimulated HK*-fbp1Δcig1*Δ cohort showed lower levels of CD4^+^ IFNγ producing cells compared to HK*-fbp1*. To further confirm HK-*fbp1* primed CD4^+^ T cells response to Cig1 antigen, we conducted T cell recall experiments and confirmed Cig1 antigen can elicit a protective cytokine response (IFNγ, IL-17A, and IL-2) in HK-*fbp1* primed T cells from the mLN whereas HK-*fbp1Δ cig1*Δ primed CD4^+^ T cells did not (**Figure 6C-6D**).

**Figure 6:**
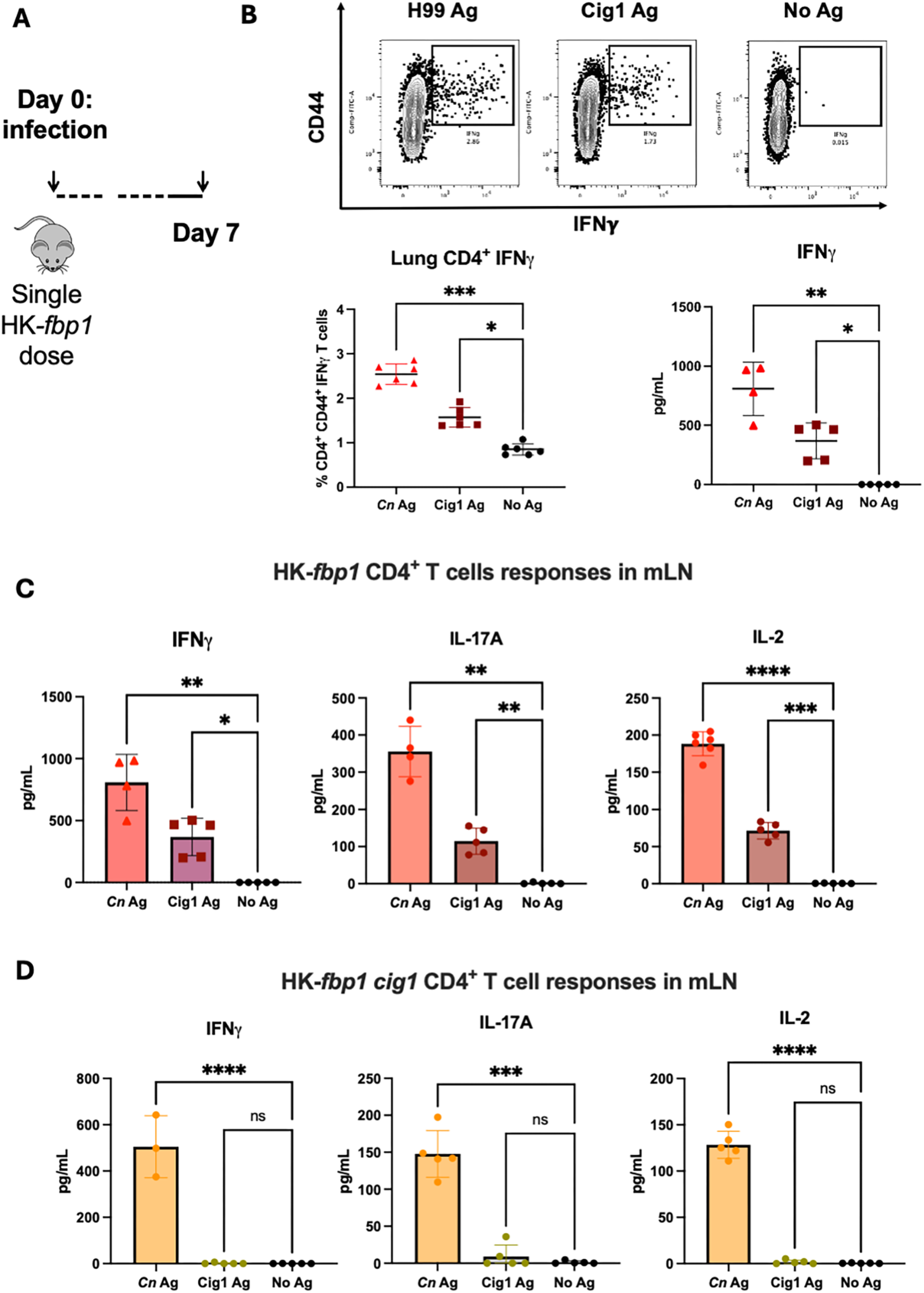
HK-*fbp1* primed lymphocytes respond to Cig1 as a stimulating immunogen. **A.** Schematic of short-term vaccination model. Balb/C mice (n=7-10) are intranasally administered one dose of 5×10^7^ HK-*fbp1* cells and sacrificed 7 days post vaccination. **B.** Top panels flow plots are representative of IFNγ producing HK-*fbp1* primed CD4^+^ T cells from whole lung suspension after ex-vivo restimulation with either whole H99 sonicated antigen, purified Cig1 antigen, or no antigen. Bottom panels show quantification results of the percentage of IFNγ producing CD4^+^ T cells and concentration of IFNγ produced by HK-*fbp1* T cells from mLN primed with *Cn* antigen or purified Cig1 antigen. **C-D.** Comparison of IFNγ, IL-17A, and IL-2 production by HK-*fbp1* (**C**) or HK-*fbp1 cig1* (**D**) primed mLN CD4^+^ T cells after ex-vivo restimulation with either whole H99 sonicated antigen, purified Cig1 antigen, or no antigen. * p<0.05, **p<0.005, ***p<0.0005, ****p<0.0001, One-way ANOVA with Dunnett’s multiple comparisons test. Data is representative of two independent experiments.

These findings demonstrate that Cig1-specific T cells are primed after challenge with *fbp1*Δ and this response is specific since immunization with *cig1*Δ failed to activate any Cig1-specific T cells. We then progressed to test the contribution of Cig1 in HK-*fbp1* vaccine protection against H99 wildtype challenge. Following our standard two dose vaccination approach, we vaccinated mouse groups with either HK-*fbp1*, HK-*fbp1Δcig1*Δ, and a no-vaccination control (**Figure 7A**). At a full dose (5×10^7^ cells/mouse) intranasal vaccination we found there was no difference in survival following H99 challenge (**Figure 7B**). As the HK-*fbp1* vaccination is dose dependent as described in previous studies^9^, we compared the two vaccinations again at half dose (2.5×10^7^ cells/mouse) to determine whether difference in protection could be distinguished. Interestingly, at half dose intranasal vaccination we discerned that HK-*fbp1Δcig1*Δ had lower protection against H99 challenge compared to HK-*fbp1 vaccination (***Figure 7C**). Additionally, we did not see a difference in lung fungal burden or dissemination to the brain in remaining survivors.

**Figure 7:**
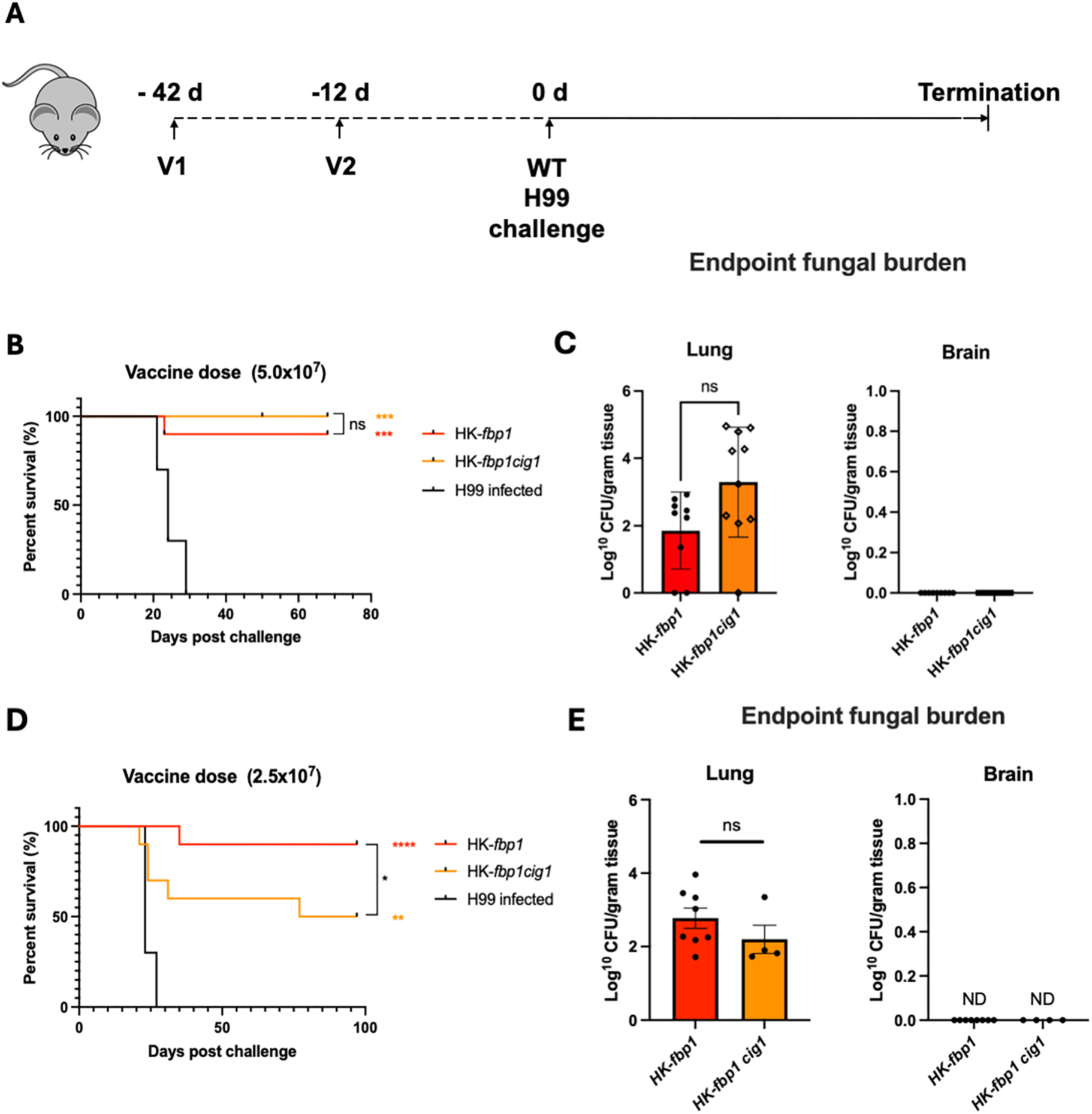
Cig1 plays a role in HK-*fbp1* protection observed only at half dose vaccination. **A.** Schematic of vaccination strategy. Mice are vaccinated with HK-*fbp1* or HK-*fbp1cig1* strains twice before challenged with 1×10^4^ H99 wildtype strain. **B-C.** Survival rate of mice vaccinated with HK-*fbp1* or HK-*fbp1* cig1 strains at full dose (5x 10^7^ cells/mouse) and challenged with H99 (B) and the fungal burden of remaining survival animal at the endpoint of the experiment (C). (D-E) Survival rate of mice vaccinated with HK-*fbp1* or HK-*fbp1* cig1 strains at half dose (2.5x 10^7^ cells/mouse) and challenged with H99 (D) and the fungal burden of remaining survival animal at the endpoint of the experiment (E). *p<0.05, ***p<0.0005, ****p<0.0001 (log rank [Mantel-Cox] test). Data is representative of two independent experiments.

In aggregate, the results from these studies provide evidence that Cig1 is a novel Fbp1 substrate that contributes to the immunogenicity and elicited protection of HK-*fbp1* vaccination against *C. neoformans* challenge.

## Discussion

In this study, we identified for the first time a single antigen, Cig1, that alters the immunogenicity of *fbp1*Δ that contributes to the protection conferred by HK-*fbp1* vaccination against *C. neoformans* challenge. Furthermore, studies in this paper show that capsule is required but not sufficient for HK-*fbp1* vaccine protection. Identifying the importance of capsule in HK-*fbp1* builds on a growing consensus that capsule requirement for protection in several whole cell *C. neoformans* based vaccine candidates including *sgl1*Δ and Znf2^OE^ ^15,39^. While other studies have revealed loss of capsule results in diminished or complete loss of protection, this present study supports that loss of capsule directly alters the protective function of HK-*fbp1* primed T cells noted by reduced production of key inflammatory cytokine IFNγ. These findings emphasize that immunogenic antigens present on the cell surface or embedded in the capsule itself are critical for HK-*fbp1* protection, as unmasking of several antigens present in the cell wall (i.e., β-glucans, chitin/chitosan) did not enhance protection. Immunofluorescent experiments with HK-*fbp1* vaccinated sera revealing antibodies binding to distinct inner regions of the *fbp1*Δ capsule supported this hypothesis.

The potential for mannoproteins as immunogens to be utilized in vaccination approaches has been described comprehensively in previous studies^36^. Extensive mannosylation of mannoproteins is considered a primary contributor to enhanced immunogenicity and key to identification by phagocytes and stimulation of CD4^+^ T cells^9,23,24^. The role of Chitin deacetylases (Cda1, Cda2, and Cda3) in maintaining cell wall integrity by modulating chitin to chitosan is a primary example of mannoprotein importance in fungal vaccine development^14,31,34,40–42^. Although, in this Cda deficient vaccination model, the enhanced immunogenicity is due more so to the loss of chitin deacetylase activity disturbing cell wall integrity rather than the immunogenicity of the enzymes themselves^14,40^. In the current study, we have ascertained that Cig1 is an overexpressed immunogenic antigen in *fbp1*Δ that contributes to eliciting protective inflammatory cytokines from HK-*fbp1* primed CD4^+^ T cells. In wildtype parental strain H99, Cig1 is lowly expressed. This may be due to tight regulation so fungal cells can evade host immune response.

In this study we identified that cryptococcal antigen Cig1 interacts with Fbp1. While we determined there is a direct protein-protein interaction between Cig1 and Fbp1, we unexpectedly identified that loss of Fbp1 also alters transcriptional expression of Cig1 as observed in qRT-PCR data. Fbp1 is part of the SCF E3 ligase complex that transiently interacts with substrate proteins set for degradation. We recently identified CDK-related kinase (Crk1) is a downstream substrate regulated by Fbp1 via its PEST domain sequence required for ubiquitin proteolysis^18^. In the absence of Fbp1, Crk1 is accumulated and promotes the phosphorylation of Gpa1 G protein to contribute to “titan cell’ formation. Furthermore, Crk1 stabilization by deletion of its PEST domain resulted in similar phenotype to that of *fbp1*Δ including virulence attenuation in murine survival and production of IFN-γ and IL-17 by CD4^+^ T cells^18^.In our studies we found that Cig1 is also upregulated in Crk1 overexpression strains (Crk1^OE^ and Crk1^ΔPEST^ strains). Previous studies have shown that the cyclic-AMP/protein kinase A (PKA) can alter cryptococcal capsule through regulation of pH responsive transcription factor Rim101^43,44^. Meanwhile, other studies have shown evidence to support the PKA pathway can alter the Ubiquitin Proteosome System (UPS) and function of the endoplasmic reticulum^45,46^ where mannoproteins are synthesized and later shuttled to the Golgi apparatus via the secretory system^46–48^. From our results, we hypothesize a working model where regulation of Cig1 by Fbp1 occurs potentially through Crk1 and Rim101 may be a novel regulatory pathway at transcriptional level, in addition to its Fbp1-mediated post-translational regulation. Future studies focused on confirming this potential pathway would be critical to understanding how loss of Fbp1 in the UPS directly impacts the expression of Cig1.

While these findings may reveal a novel regulatory pathway between Fbp1, Cig1, Crk1, and Rim101, the impact of CIG1 deletion in the context of HK-*fbp1* vaccination is modest. The modest difference in vaccine protection between HK-*fbp1* and HK-*fbp1Δcig1*Δ vaccinated animals indicates that while Cig1 is a highly expressed immunogen in the *fbp1*Δ mutant, it is clearly not sufficient for HK-*fbp1* vaccine induced protection. The likelihood of one singular antigen bearing the responsibility of complete immunogenicity in an F-box deficient strain is low considering the complexity of fungal immunogens. We acknowledge that Fbp1 likely regulates multiple different immunogenic factors and Cig1 is one of them, which warrants further investigation.

In aggregate, our studies provide evidence that mannoprotein Cig1 and its presence embedded within the cryptococcal capsule plays an important role in shaping the immunogenicity of HK-*fbp1* vaccination induced protection against *C. neoformans* challenge. These findings mark the first identification of an immunogenic antigen that directly contributes to protection in the HK-*fbp1* vaccination model. The recent report on mRNA-based vaccine using Cig1 as an antigen further confirms the immunogenicity of Cig1 as an important immunogen in *C. neoformans*^38^. Furthermore, we have identified Fbp1 as potential novel regulator of Cig1 through modulation of downstream substrates but remains to be further characterized. In conclusion, our data build on the growing body of knowledge of the relevance of mannoproteins as key targets for fungal vaccine development that may also play a role in other current fungal vaccine candidates.

## Methods

### Animals and fungal cultures

Female mice with an average weight of 19 to 23 g were used throughout these studies. BALB/c, and C57BL/6 genetic background of mice were purchased from the Jackson Laboratories. Animal studies were performed at the Public Health Research Institute Animal facility. All studies were conducted by following biosafety level 2 (BSL-2) protocols and procedures approved by the Institutional Animal Care and Use Committee (IACUC) and Institutional Biosafety Committee of Rutgers University. *Cryptococcus neoformans* clinical strain H99 and its mutants were cultured on yeast extract-peptone-dextrose (YPD) medium.

### qRT-PCR

*Cryptococcus* cells from overnight culture were collected and washed two times with ddH_2_O, frozen down to -80°C for 6-8 hours and lyophilized to powder form overnight. Total RNA extraction and first-strand cDNA synthesis were performed as described previously^18^. Expression of *CIG1, CMP1, CFO1, MP88, MP84, MP98, MP115* and *GAPDH* were analyzed using SYBRGREEN advantage QPCR premix reagents (Takara). Gene expression levels were normalized using the endogenous control gene *GAPDH*, and the relative levels were determined using the comparative threshold cycle (*C_T_*) method. Quantitative real-time PCRs (qRT-PCRs) were performed using Aria MX Real Time PCR System (Agilent Technologies).

### Macrophage phagocytosis assays

Phagocytosis assays were performed in 96-well plates, using J774 macrophages at a concentration of 2.5x 10^4^ cells/well that were allowed to double overnight at 37°C in activating medium (DME medium, 50 U/ml of gamma interferon [IFN-], 1 g/ml of lipopolysaccharide [LPS]). Phosphate buffered saline pH 7.4 (PBS) washed fungal cells were opsonized with 20% mouse complement (Pel-Freez, AK) and added to the macrophages at an effector-to-target ratio of 1:2. Phagocytosis was allowed to occur for 2 h at 37°C in 10% CO_2_. The cells were then washed three times with 1X PBS and fixed with methanol at 4°C for 30 min. Giemsa stain was added to the wells at a dilution of 1:20, and the plates were incubated at room temperature for 30 min. The wells were washed once with 1X PBS and analyzed using an inverted microscope. For each well, 3 different fields were counted, for a total of at least 100 macrophages. The percent phagocytosis was determined by dividing the number of macrophages that contained *C. neoformans* by the total number of macrophages counted.

### Generation of tagged protein strains

The *CIG1* full-length gDNA was amplified with primers CX2373 and CX2375. The *CIG1* fragment was cloned into the BamHI/ SpeI sites of a vector pCXU254 via In-Fusion HD cloning kit (Takara) This cloning produced plasmid pCXU484 which contains a *CIG1*:mcherry fusion that is under the actin promotor. The above plasmids were biolistically transformed into the H99 strain to generate strains CUX1454.

### Antigen pulldown assays

The Cig1 protein from Cig1:mCherry total protein lysate was pulled down by using Protein G Dynabeads (Invitrogen) bound to rat anti-mCherry monoclonal antibody (Invitrogen) and eluted with 0.1 M Glycine pH 2.0 and neutralized with 1.0M Tris HCL (pH 8.0). The purified Cig1:mCherry protein was analyzed by immunoblotting with anti-mCherry (1:2000).

### Co-immunoprecipitation assay

Total proteins were purified and analyzed by immunoblotting with rabbit anti-FLAG (GeneScript) and rat anti-mCherry (Invitrogen) monoclonal antibodies. Proteins were pulled down by using anti-FLAG affinity gel (Sigma, A2220) incubated overnight and run on 12% SDS-PAGE gel. Samples were analyzed by immunoblotting with anti-FLAG antibody to detect Fbp1:FLAG or with anti-mCherry antibody to detect Cig1:mCherry.

### In vivo Ubiquitination assay

CUX1454 (P_ACT1_-Cig1:mCherry) was cultured in YPD at a set OD_600_ 1.0 overnight. Cells were collected, washed and cultured in YPD in the presence or absence of 20 µM MG132 for 6 h. Total proteins were extracted in lysis buffer (50 mM Tris-HCl, 150 mM NaCl, 1% Triton X-100, 1 mM EDTA, 10 µM PMSF and 1×EDTA-free protease inhibitor). Accumulation of Cig1 polyubiquitination (Cig1-(Ubn) was detected using anti-mCherry antibody (Invitrogen, M11217, 1:2000)

### Vaccination strategy

*C. neoformans* mutant strains were heat killed by following a previously described procedure^9^. Briefly, fungal cells from YPD overnight cultures were spun down and washed twice with sterile 1X PBS and resuspended and heated on a hot plate at 75°C for 105 min, vortexed every 15 minutes to ensure proper killing of all cells. To ensure killing of cells via heat treatment, the heat killed cell suspension was plated on YPD agar plates and incubated at 30°C for 2 days. Mice were vaccinated intranasally in a 50 μL volume of either with 5×10^7^ or 2.5×10^7^ heat-killed fungal cells after being anesthetized with a mix of ketamine (12.5 mg/ml) and xylazine (1 mg/ml) at day -42. Each group of 8 to 10 mice was boosted again with the same dose of heat-killed fungal strains at day -12. A group of unvaccinated mice were utilized as a control. Vaccinated and unvaccinated cohorts were challenged with 1×10^4^ live H99 cells via intranasal inoculation or other fungal species as specified below. Infected animals were weighed and monitored daily for disease progression, and mice displaying severe disease symptoms were euthanized. All survivors were euthanized on day 65-100 after challenge with live H99 cells unless otherwise specified. Survival data from the murine experiments were statistically analyzed between paired groups by using the log rank (Mantel-Cox) test with PRISM version 10.0 (GraphPad Software, San Diego, CA) (P <0.05 were considered statistically significant). The resulting data were plotted against time.

### Lung processing

Single-cell suspensions of pulmonary cells were prepared for flow cytometric analysis according to previously described methods^17^. Briefly, lung tissue was minced in 5 ml of 1X PBS containing 3 mg/ml collagenase type IV (Worthington). Samples were incubated at 37°C for 45 min and washed with 1X PBS three times. After digestion, residual RBCs were removed using RBC lysis buffer (155 mM NH_4_Cl and 10 mM NaHCO_3_, PH 7.2). Total numbers of lung cells collected for each sample were determined by counting numbers of cells on a hemocytometer. Lung single-cell suspensions were stained for CD4^+^ T cells (CD45 [30-F11 BUV395] and CD4^+^ [ CD4 RM4-5, BV421]), and CD8^+^ T cells [CD8α 53-6.7 BV711]. All antibodies used for lung staining were from BD Biosciences. All samples were analyzed using a BD LSR Fortessa flow cytometer and FlowJo software.

### Intracellular cytokine staining of T cells harvested in BALF and flow cytometry

For analyzing host immune responses, BALF samples were harvested at the endpoint after inoculation and prepared for flow cytometry analysis as previously described^17^. Briefly, BALF was collected in 5 ml of 1x PBS buffer using a catheter inserted into the trachea of animal post euthanasia. Reb blood cells (RBC) were removed using RBC lysis buffer. BALF cells were then plated in a 96-well round-bottom plate and restimulated using BD-leukocyte activation cocktail containing BD GolgiPlug (BD Biosciences) according to the manufacturer’s instructions. Six hours after activation, BALF cells were surface stained with fluorescently labeled antibodies against Thy1.2, CD4, and CD8. Samples were fixed in 1% paraformaldehyde overnight. Prior to intracellular staining, the samples were permeabilized with 1X BD Perm/Wash buffer according to the manufacturer’s instructions. Intracellular cytokine staining (ICCS) was done using fluorescently labeled antibodies against IFN-γ, IL-17A, TNF-α, and IL-13 diluted in 1X BD Perm/Wash for 30 min on ice. Cells were cell surface stained for T cells with Thy1.2 (53-2.1 PE-Cy7), CD4 (RM4-5 BV421), CD8 (53-6.7 BV711) and ICCS for IFN-γ (XMG1.2 PE), IL-17A (eBio17B7 APC), TNF-α (MP6-XT22 Alexa Flour 700), and IL-13 (eBio13A FITC).

### Lung lymphocyte ex vivo restimulation

Lung samples were minced in 5 ml of 1X PBS containing 3 mg/ml collagenase type IV (Worthington) and incubated at 37°C for 45 min. Samples were then washed with 1X PBS and then resuspended in 40% Percoll (Sigma) layered on top of 66% Percoll and centrifuged at 2000 rpm for 20 min to purify lung immune cells. Single cell lung suspension samples of single dose vaccinated mice were pooled together (n=10) incubated on a 48 well plate with CD28 1:100 dilution (Biolegend clone 37.51) and antigen for 2 hours incubated at 37°C with 5% CO_2_. After restimulation with antigen, samples were incubated with 1X Brefeldin A (Invitrogen) for 4 hours at 37°C with 5% CO_2_. Lung lymphocytes were then stained and fixed for overnight intracellular cytokine staining with Foxp3 fix/permeabilization kit (eBioscience) and later analyzed via flow cytometry. Activated CD4^+^ T cells production of cytokines was gated as near-IR LIVE/DEAD (Invitrogen)^-^, CD90^+^, NK1.1^-^, Ly6C^-^,Ly6G^-^,SiglecF^-^,CD11c^-^,TCRβ^+^,TCRγδ^-^,CD8^-^,CD4^+^, and CD44^+^.

### CD4^+^ T cell isolation and CD4^+^ T cell recall response

Lung-draining mediastinal lymph nodes (mLNs) were collected and placed in 10 ml of 1X PBS. Total lymphocyte cell suspensions were prepared by gently releasing the cells into the 1X PBS by grating the lymph nodes with the frosted ends of two glass slides. Individual samples from each group were pooled (5 to 6 mice). CD4^+^ T cells were purified using a negative-sorting CD4^+^ isolation kit (Miltenyi Biotec, Inc., Auburn, CA). CD4^+^ T cell isolation was done by following the manufacturer’s instructions and were consistently found to be (90% pure, as assessed by flow cytometry. Purified CD4^+^ T cells (2 x 10^5^) were cultured with T cell-depleted antigen-presenting cells (3 x 10^5^) in RPMI containing 10% fetal calf serum (FCS) and penicillin-streptomycin (2,200 U/ml; Gibco). The cultures were plated in flat-bottom 96-well plates and incubated at 37°C with 5% CO2 for 72 hr. Antigen-presenting cells were prepared from the spleen of syngeneic, uninfected donor mice. In brief, splenic cell suspensions were depleted of T cells by antibody complement-mediated lysis. Splenic cells were incubated with anti-Thy1.2 antibodies and rabbit complement (Low Tox; Cedarlane Labs, Hornby, ON, Canada) at 37°C for 60 min. To measure Cryptococcus-specific responses, CD4^+^ antigen-presenting cell cultures were incubated with sonicated H99 as a source of fungal antigens. The amount of antigen used was adjusted to a multiplicity of infection of 1:1.5 (antigen-presenting cell:yeast). The fungal growth inhibitor voriconazole was used at a final concentration of 0.5 mg/ml to prevent any fungal cell outgrowth during the culture period. After 72 h post culture, with initiation at 37°C with 5% CO_2_, supernatants were collected for cytokine analysis by enzyme-linked immunosorbent assay (IL-2, IL-5, IL-17, TNF-α,and IFN-γ, Invitrogen) by following the manufacturers’ instructions.

### Fungal burdens in infected organs

Infected animals were sacrificed at designated time points and the endpoint of the experiment according to the Rutgers Institutional Animal Care and Use Committee (IACUC)-approved animal protocol. Infected lungs, brains, and spleens were also isolated and homogenized in 3 mL of 1X PBS. Resuspensions were diluted, 200 uL of each dilution was spread on YPD medium with ampicillin and chloramphenicol, and numbers of colonies were determined after 3 days of incubation.

### Statistics

GraphPad Prism version 10.2.2 (GraphPad Software, La Jolla, CA) was used for statistical analyses and graph design. Survival data from the murine experiments were statistically analyzed between paired groups by using the log rank (Mantel-Cox) test where *P* values of <0.05 were considered statistically significant. Statistical analysis of in vivo and in vitro experiments with three or more groups, Shapiro-Wilk test for normality and Brown-Forsythe test for equal variance amongst groups were performed. For multiple-comparison analysis of three or more groups that met normality and equal variance conditions, a One-Way ANOVA with Dunnett’s multiple comparisons test or Kruskal-Wallis test with Dunn’s multiple comparison test was used.

### Ethics statement

The animal studies described here were compliant with all the provisions established by the Animal Welfare Act and the Public Health Services Policy on the Humane Care and Use of Laboratory Animals. All studies were performed in accordance with study protocols reviewed and approved by the IACUC of the Rutgers University Newark campus.

## Acknowledgments

We thank James Kronstad for generously providing the Cig1 mutant strains. We thank formal lab member Chengjun Cao for sharing unpublished data utilized to identify Cig1 as a potential substrate. Additionally, we thank Darin Weisner for sharing reagents, materials, and intellectual support to these studies. We also thank Jason Weinstein for helpful discussions to better these studies. This study is supported by NIH grants AI169769 and AI141368 to AR and CX. The Xue lab is also supported by NIH grants AI123315 and AI155647. AR holds an Investigators in the Pathogenesis of Infectious Disease Award from the Burroughs Welcome Fund.

