## Supplemental materials for "Mannoprotein Cig1 Contributes to the Immunogenicity of a Heat-Killed F-box Protein Fbp1 *Cryptococcus neoformans* Vaccine Model"

### Supplemental figures

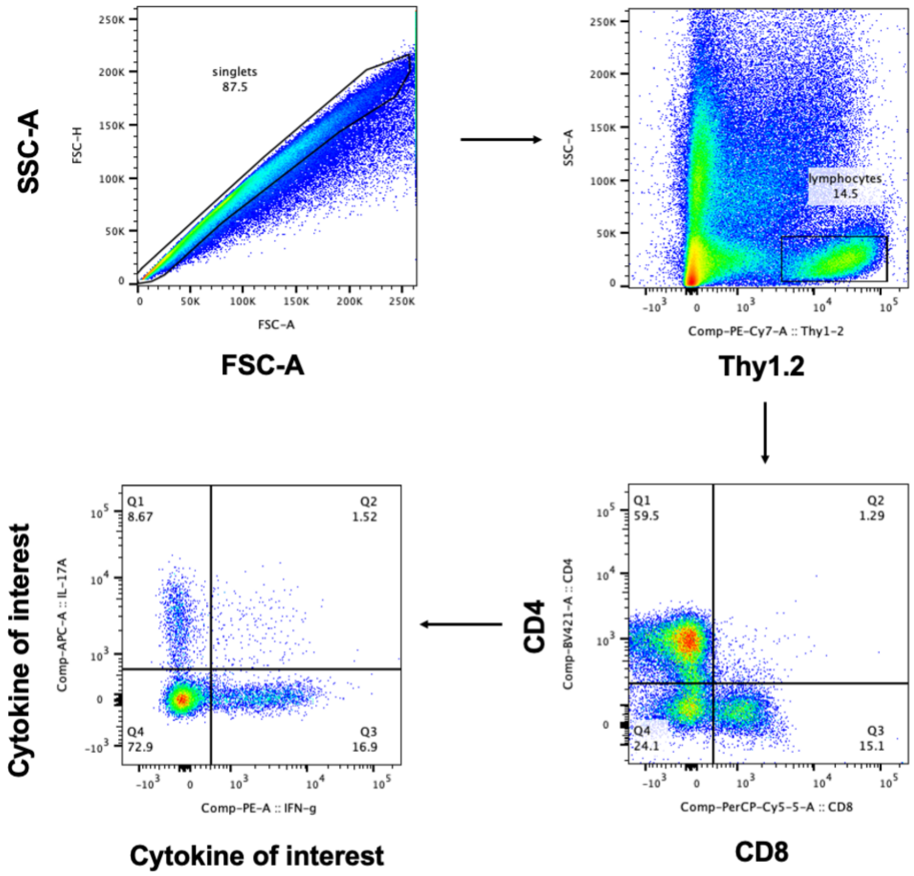


**Figure S1. Bronchoalveolar lavage fluid (BALF) CD4⁺ T cell isolation gating strategy.**

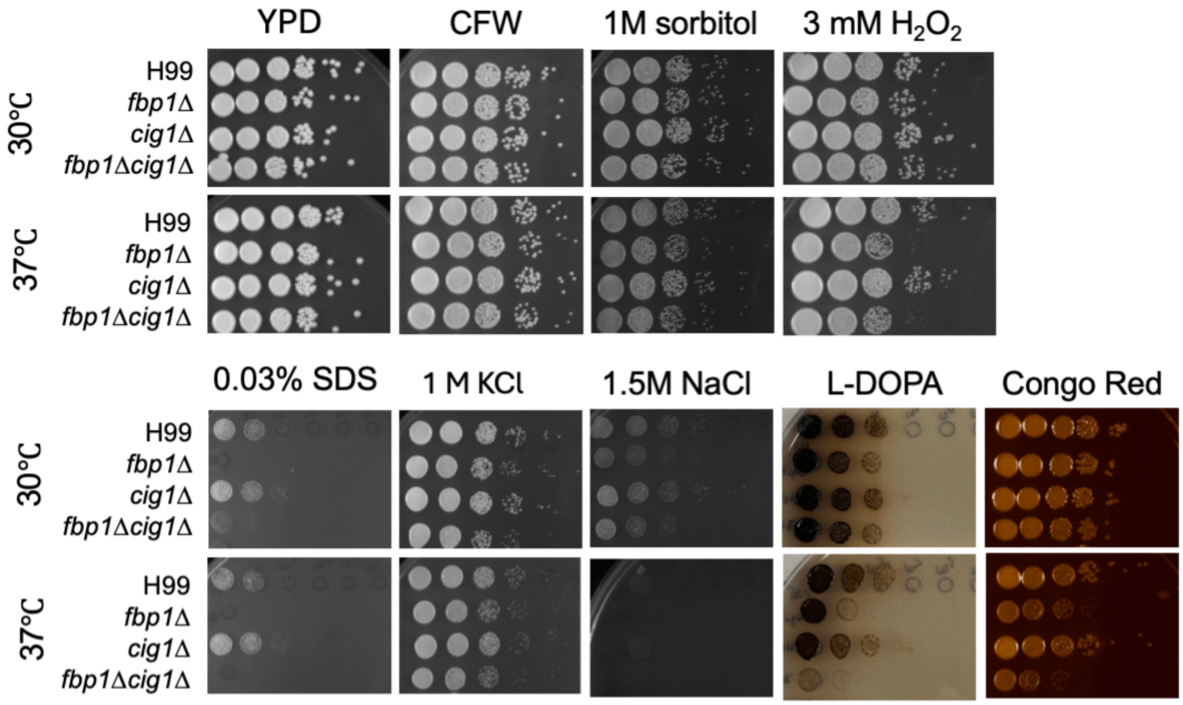


**Figure S2. Cig1 mannoprotein is upregulated in *fbp1*Δ and alters *Cn* cell wall integrity.** Comparison of parental H99, *fbp1*Δ***,*** *cig1*Δ***,*** and *fbp1*Δ *cig1*Δ mutant strains on different growth conditions at 30°C and 37°C.

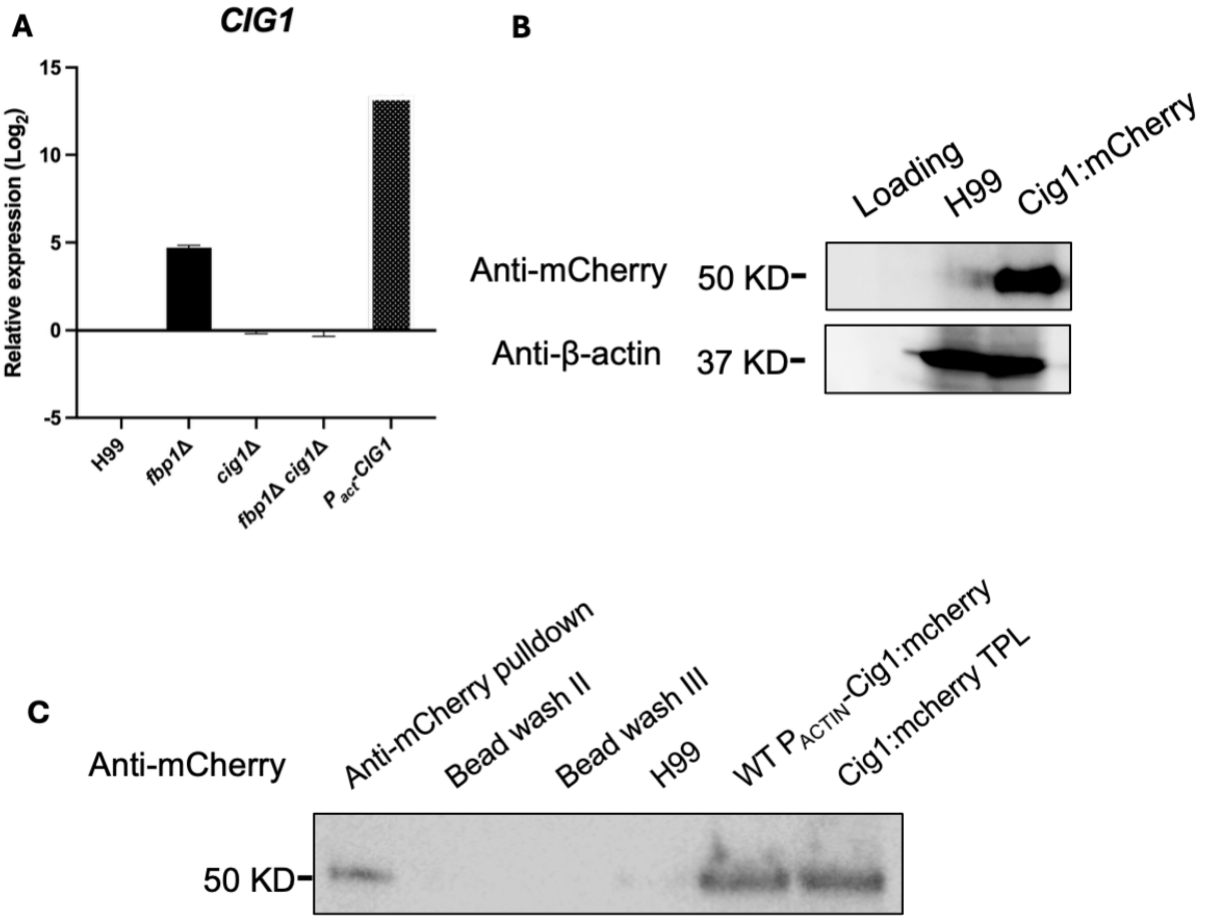


**Figure S3. Construction and validation of *$P_{ACTIN}$-CIG1:mcherry* strain for Cig1 protein enrichment. A**. qRT-PCR of Cig1 expression in mutant strains. **B.** Western blot analysis of Cig1-mCherry tagged protein with β-actin as a control. **C.** Protein enrichment of Cig1 from *$P_{ACTIN}$-CIG1:mcherry* expressing *C. neoformans* total protein lysate via co-IP with rat anti-mCherry antibody bound to protein G Dynabeads (Invitrogen).

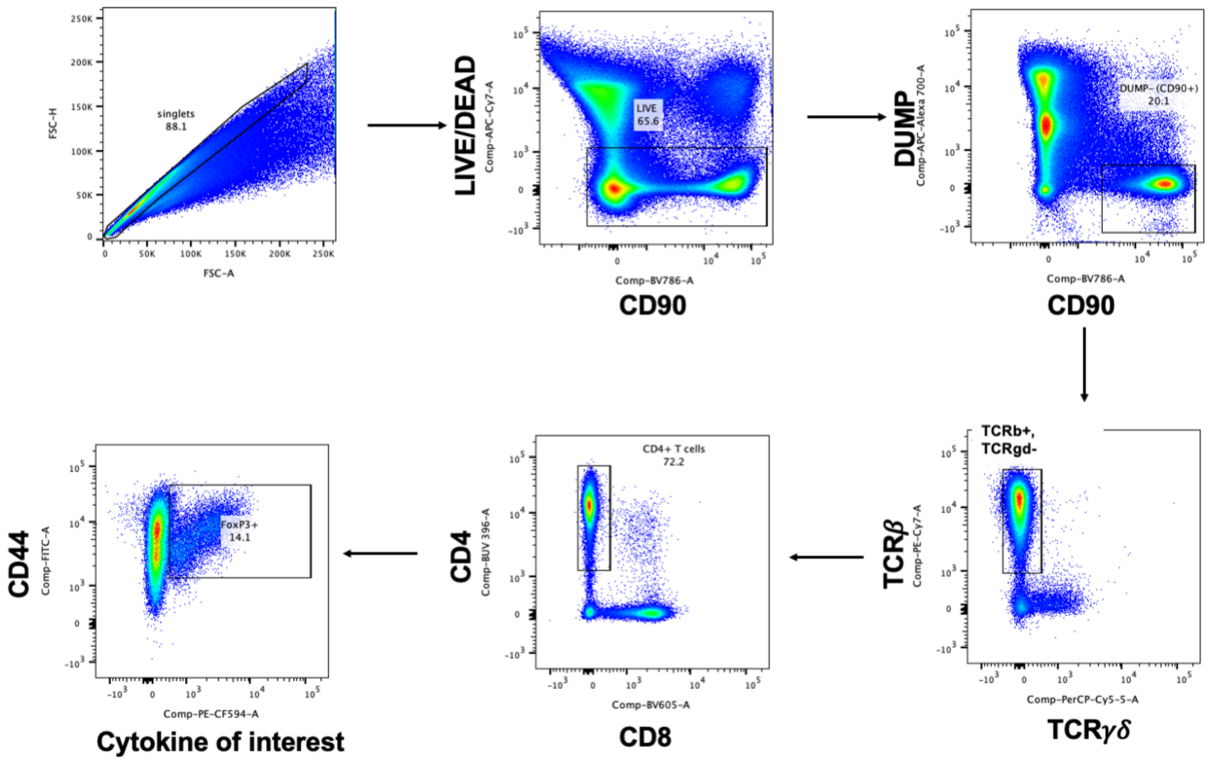


**Figure S4. CD4⁺ T cell purification gating strategy in ex vivo antigen restimulation experiment.**

**Supplementary table 1: List of strains and plasmids used in this study.**

| **strains** | **genotype** | **Source/reference** |
|---|---|---|
| H99 | MATα | Bjorklund et al 2019 |
| KN99a | MAT**a** | Lloyd et al 2013 |
| CUX2 | MATα *fbp1Δ::NEO* | Cao et al 2022 |
| CUX1446 | MATα *cig1Δ::NAT* | Kronstad |
| CUX1454 | MATα $P_{ACT1}$*-CIG1:mCherry-NAT* | This study |
| CUX1455 | MATα *fbp1Δ::NEO cig1Δ::NAT* | This study |
| CUX1459 | MATα *fbp1Δ::NEO ura5* $P_{ACT1}$*-FBP1:FLAG-URA5* $P_{ACT1}$*-CIG1:mCherry-NAT* | This study |
| CUX1291 | MATα $P_{ACT1}$*-CRK1:mCherry-NAT* | Cao et al 2022 |
| CUX1293 | MATα $P_{ACT1}$*-CRK1*$^{\Delta PEST}$*:mCherry-NAT* | Cao et al 2022 |
| **Plasmids** | **Description** | **Source/reference** |
| pCXU254 | $P_{ACT1}$-*CHO1*:mCherry-*NAT* | Konarzewska et al 2019 |
| pCXU484 | $P_{ACT1}$*-CIG1:mCherry-NAT* | This study |

**Supplementary table 2: List of primers used in this study.**

| Primers | Sequences (5'-3') | References/Note |
|---|---|---|
| CX49 | TGAGAAGGACCCTGCCAACA | GAPDH F qPCR |
| CX50 | ACTCCGGCTTGTAGGCATCAA | GAPDH R qPCR |
| CX18993 | GACGACATCTATAGCAGATC | MAT alpha F |
| CX18994 | CCAAAAGCTGATGCTGTGGA | MAT alpha R |
| CX18995 | TCCACTGGCAACCCTGCGAG | MAT **a F** |
| CX18996 | ATCAGAGACAGAGGAGCAAGAC | MAT **a R** |
| CX284 | GGAATCAACGAAGATCTGAAGG | FBP1 F positive |
| CX1104 | TGTGGATGCTGGCGGAGGATA | FBP1 R positive |
| CX282 | GACCGTCAAGAAGCAACTCG | FBP1 F negative |
| CX283 | CAGATTGATAGCTTGCAGCTTC | FBP1 R negative |
| CX2259 | CCTCGTGGTTTCTCAGCTTC | CIG1 F qPCR |
| CX2260 | CCACCGCTCATAGGGTAAGA | CIG1 R qPCR |
| CX2261 | ACATGACCGCCCTCAGTAAC | MP98 F qPCR |
| CX2262 | CCTCGGCAATCATACGAACT | MP98 R qPCR |
| CX2263 | TTCGAGGCTGGAAAGTCCTA | CFO1 F qPCR |
| CX2264 | ACTTCGACACCGTCCATTTC | CFO1 R qPCR |
| CX2265 | ACCTACCATGCTTCCCCTCT | CMP1 F qPCR |
| CX2266 | GTCGGAGGACTGGTTGACAT | CMP1 R qPCR |
| CX2267 | GAATGCCAGCACACTCTTGA | MP88 F qPCR |
| CX2268 | GCTGCTGGAAGGTAGAGGTG | MP88 R qPCR |
| CX2269 | GCGTTGAGGCTTACTTCGAC | MP84 F qPCR |
| CX2270 | GTTAAAAGCATCGGCCACAT | MP84 R qPCR |
| CX2271 | CGCCTTGAGTGCAGACAATA | MP115 F qPCR |
| CX2272 | TGTTGCGGGAAAGGATAGTC | MP115 R qPCR |
| CX2373 | CAACATGTCTGGATCCATGATTTTTAATCGTTTCACATTCA | CIG1 F infusion pCXU254/BamHI |
| CX2375 | CCATTCTAGAACTAGTGAGACGCTCCTTGGTGGGT | CIG1 R infusion pCXU254/SpeI |
